# Reconstructing physical cell interaction networks from single-cell data using Neighbor-seq

**DOI:** 10.1101/2022.04.15.488517

**Authors:** Bassel Ghaddar, Subhajyoti De

## Abstract

Cell-cell interactions are the fundamental building blocks of tissue organization and multicellular life. We developed Neighbor-seq, a method to identify and annotate the architecture of direct cell-cell interactions and relevant ligand-receptor signaling from the undissociated cell fractions in massively parallel single cell sequencing data. Neighbor-seq accurately identifies microanatomical features of diverse tissue types such as the small intestinal epithelium, terminal respiratory tract, and splenic white pulp. It also captures the differing topologies of cancer-immune-stromal cell communications in pancreatic and skin tumors, which are consistent with the patterns observed in spatial transcriptomic data. Neighbor-seq is fast and scalable. It draws inferences from routine single-cell data and does not require prior knowledge about sample cell-types or multiplets. Neighbor-seq provides a framework to study the organ-level cellular interactome in health and disease, bridging the gap between single-cell and spatial transcriptomics.

## INTRODUCTION

The spatial context of cells in a tissue and their resulting cell-cell communications influence numerous processes, including cellular differentiation, organ development and homeostasis, and immune interactions in disease (1). Few high-throughput methods exist that can resolve direct cellular communications *in vivo* at single-cell resolution. Single-cell RNA sequencing (scRNA-seq) can identify cell-types and states in heterogeneous tissues, but the tissue structure is largely destroyed in the process (2). Microscopy-based methods such as RNAscope and FISH can interrogate only preselected genes at high spatial resolution. Spatial transcriptomics allows profiling of microscopic regions, but still samples 10-100 cells per region (3, 4). Sequencing of partially dissociated tissues and subsequent multiplet deconvolution permits inference of physical cell interactions, but this requires specialized experimental modifications (5–8). A general method that can infer direct cell-cell interactions and concurrent transcriptomic changes *in vivo* at single cell resolution would provide unprecedented insight into the building blocks of tissue architecture in healthy and diseased tissues.

Cell aggregates (multiplets) naturally arise in scRNA-seq experiments when two or more cells are captured in the same reaction droplet, and they typically represent at least several percent of all capture events (9, 10). Such multiplets occur primarily due to incomplete tissue dissociation (6), or occasionally by random co-encapsulation. We developed Neighbor-seq, a method to infer physical cell-cell communications by identifying, annotating, and analyzing cell multiplets from the undissociated cell fractions in scRNA-seq data using computational approaches (see Methods, **Fig. 1A**). Neighbor-seq provides a framework to study the cellular interactome in health and disease using standard scRNA-seq data.

**Figure 1.**
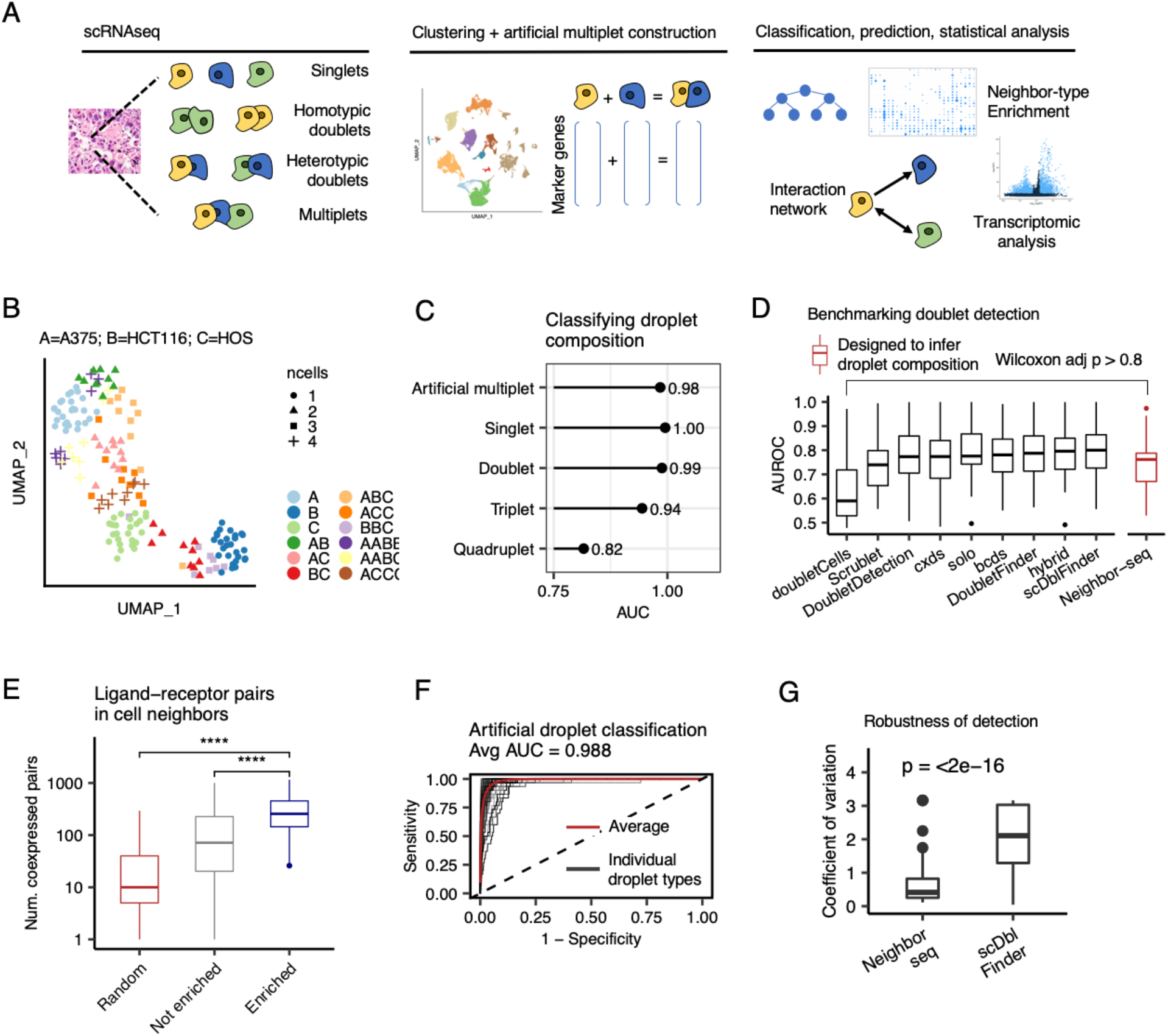
Benchmarking cell neighbor detection and annotation. **(A)** A schematic representation of the Neighbor-seq workflow. **(B)** Uniform manifold approximation and projection (UMAP) of barcode RNA sequencing data singlets and multiplets of known composition from 3 cancer cell lines, colored by cell-type identities and shaped by the number of cells per barcode. **(C)** Neighbor-seq barcode composition annotation performance of cell-line barcodes in (B), plotted by known barcode type. AUC = area under the receiver operator curve. **(D)** Benchmarking Neighbor-seq doublet detection against 9 other methods using 16 datasets of diverse tissue types with experimentally annotated doublets (see **Table S1**). Among these methods, only Neighbor-seq is explicitly optimized to infer doublet composition. Comparison of singlet vs. doublet classification area under the receiver operator curve (AUROC) distributions are shown. Boxplots show median (line), 25^th^ and 75^th^ percentiles (box) and 1.5xIQR (whiskers). Points represent outliers. **(E)** Comparison of the number of co-expressed ligand-receptor pairs in enriched (statistically significant) doublet types, not enriched doublets, and random synthetic doublets from the benchmarking studies in (D). Descriptions of the boxplots are as in (D). (Wilcoxon tests, ****p<0.0001. See also **Fig. S1A**). **(F)** Neighbor-seq example receiver operator curve for classifying artificial multiplet types in the cline-ch study, one of the benchmark studies used in (D). AUC, area under the curve. **(G)** Robustness of cell neighbor type annotation. Comparison of the distribution of coefficients of variation for neighbor type counts from the cline-ch benchmark dataset across n=10 runs of Neighbor-seq and scDblFinder (Wilcoxon test, boxplots as in (D)).

## METHODS AND MATERIALS

### Neighbor-seq algorithm

Neighbor-seq is a method to infer physical cell-cell communications by identifying, annotating, and analyzing cell multiplets from the undissociated cell fractions in scRNA-seq data using machine learning approaches. The Neighbor-seq algorithm consists of the following components, each further described below: (i) barcode clustering and marker gene identification, (ii) Random Forest classifier training to identify multiplets and their cell type compositions, (iii) calculating enrichment scores for cell-cell interactions, and (iv) construction of cell-cell interactome network and analysis of cell-neighbor transcriptomes, including ligand-receptor interactions (**Fig. S4**).

#### Input and cell type identification

The input for Neighbor-seq is a cell by gene counts matrix, and optionally, cell-type cluster labels. If cell-type labels are not provided, Neighbor-seq utilizes a wrapper function to run Seurat (11) functions that TP10K normalize and scale the scRNA-seq data, find a default of 5000 variable genes, perform principle component analysis with n=50 components, and identify cell type clusters. Cell-type marker genes are then identified using the FindAllMarkers function using a default of 200 cells subsampled for each cluster, an average log fold change threshold cutoff of 1, and minimum fraction of cells expressing a gene of 0.2 for computational efficiency. If cell-type labels are known a priori, only normalization and marker finding functions are run.

#### Random Forest classifier training

Next, all homotypic and heterotypic combinations for a default of 2 cell-types are enumerated (e.g. AA, AB, etc.). Cells forming a doublet or multiplet are hereon referred to as “neighbors”, and the “neighbor-type” indicates the identities of the 2+ cells on a single barcode. For each neighbor-type, artificial multiplets are created by randomly sampling cells from the constituent cell types and the prepared input gene by cell matrix and summing their gene counts. A random forest is then trained using a balanced set of singlets and artificial multiplets to predict the cell type composition of the barcodes from the assembled dataset. Although the majority of barcodes are expected to contain single cells, a balanced training set is used to increase random forest prediction accuracy. As such, a default of 100 artificial multiplets is created for each neighbor-type, and these are pooled with a default of 100 singlets from each originally identified cell-type to create the training set. A random forest trained, and hold-out artificial multiplets data are used to assess random forest performance. The multiROC R package is used to compute receiver operator curves. The process is iterated multiple times to retrain and generate an ensemble annotation result. The trained random forest is then used to predict the barcode composition of all barcodes in the original dataset. Two random forest implementations are incorporated in Neighbor-seq: the randomForest(12) R package, run with a default of 500 trees, and a faster implementation via the xgboost(13) R package, configured with the following parameters: objective=multi:softprob, eval_metric=mlogloss, nround=1, max_depth=20, num_parallel_tree=200, subsample=0.632, colsample_bytree=sqrt(number cells)/total number of cells, and colsample_bynode=sqrt(number of cells)/total number of cell. The faster implementation is preferred with large datasets and is set as the default in Neighbor-seq. Both implementations generated consistent results on benchmark analyses (**Fig. S5b**).

#### Determining significantly enriched cell-cell interactions

Neighbor-seq calculates an enrichment score for each cell-type interaction and compares it to the distribution of enrichment scores expected by chance. The interaction enrichment score reflects the proportion of counts of a neighbor type relative to the product of the total number of edges detected from each constituent cell and all other cell types. For neighbor type *C*_*1*_*C*_*n*_ composed of cell types *C*_*1*_ … *C*_*n*_, the enrichment score is specifically defined as:

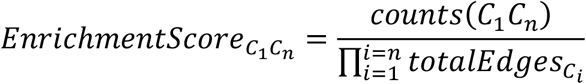

The observed enrichment score for a neighbor type is compared to that expected by chance. The null hypothesis assumes that multiplet formation is random and thus the distribution of neighbor-types follows the underlying singlet population counts. As such, for each sample in a dataset, given *n* predicted singlets and *m* predicted multiplets consisting of *x* constituent cells, Neighbor-seq simulates the synthetic creation of *m* multiplets drawing without replacement from *n+x* cells. The resulting neighbor-types are tallied and their enrichment scores are computed. This simulation is repeated for a default of 100 times; for each neighbor type, lower tailed Wilcoxon testing quantifies the probability that the simulated enrichment scores have a central tendency greater than the observed enrichment score. All probabilities are adjusted using the Holm correction.

#### Construction of a cell-cell interactome network and transcriptomic analysis

The primary outputs of Neighbor-seq are:(1) the artificial multiplet training and test sets, (2) the trained random forest, (3) the barcode classification probabilities for all barcode classes, and (4) the neighbor enrichment analysis. The latter reports the counts of each multiplet class detected in a sample as well as its enrichment score and corresponding p-value. When an ensemble result is generated by retraining the algorithm over multiple iterations, Neighbor-seq additionally reports the mean counts for each multiplet class and a combined p-value using Fisher’s method. Using the cell neighbor type enrichment data, we use igraph(14) to plot a network, and it is possible to calculate network statistics such as degree and centrality. We use ligand-receptor data from Ramilowski et al.(15) or Shao et al. (16) to identify highly expressed ligand-receptor pairs in cell neighbors. Differentially expressed genes and other transcriptomic features in neighbor-types can be assessed using Seurat(11).

### Benchmarking, robustness, and reproducibility

We used a number of different metrics for benchmarking Neighbor-seq.

#### Comparative assessment with other doublet finding methods

We obtained scRNA-seq data for the benchmark annotated doublet datasets from Xi et al.(10) (see **Table S1** for dataset details) and compared Neighbor-seq’s ability to identify singlets vs. doublets against the following 9 published methods: doubletCells(17), Scrublet(9), DoubletDetection (https://github.com/JonathanShor/DoubletDetection), cxds(18),bcds(18), hybrid(18), solo(19), DoubletFinder(20), and scDblFinder (https://github.com/plger/scDblFinder). These methods are optimized to distinguish singlets and filter out doublets in scRNA-seq data, and only scDblFinder has the ability to propose doublet composition. These methods generally rely on simulating artificial doublets, projecting data into a lower dimensional space, and using a machine or deep learning classifier to annotate barcodes. In contrast, Neighbor-seq utilizes cell-type specific clustering and marker gene sets to train a classifier directly on the gene counts of all possible expected barcode compositions, and it explicitly assigns multiplet class probabilities for each barcode rather than annotating the broader label of singlet or doublet. To evaluate Neighbor-seq, we ran Neighbor-seq using the default parameters on each benchmark dataset. To compute the probability of each barcode being either a singlet or doublet, we summed the Neighbor-seq probabilities for all doublet classes. Classification performance (area under the receiver operator curves) for the 9 benchmark doublet finding methods tested on these datasets were obtained from Xi et al.(10), who ran these methods using their default settings. Area under the receiver operator curves for the Neighbor-seq singlet vs. doublet classification were calculated as above with multiROC and compared with all other methods.

To compare the stability of doublet population predictions across multiple runs from Neighbor-seq and scDblFinder, we ran each algorithm 10 times on the cline-ch benchmark dataset obtained from Xi et al.(10) For each doublet class, the coefficient of variation of its counts (mean/standard deviation) was computed, and the distributions of coefficients of variations were compared using Wilcoxon testing.

#### Ensemble prediction

To increase reproducibility of results, artificial multiplet construction and/or random forest training can be run for multiple iterations and an ensemble results can be computed. For each neighboring cell-type pair, the mean observed counts and Fisher’s combined adjusted p-value are reported. We observed a high degree of reproducibility across multiple iterations and with a range of random seeds. These multiplet type counts and p-value thresholds for determining significant interactions were set to counts>10 and p-value<0.05 for most analyses in this study, but they can be adjusted and interpreted in light of data quality and known biology.

#### Evaluation of Ligand-receptor co-expression in cell neighbors

Ligand-receptor data was obtained from Ramilowski et al. (15) or Shao et al. (16). We compared three classes of barcodes:

(1) multiplet-classes with statistically significant enrichment, (2) multiplet-classes without statistical enrichment, and (3) randomly synthesized multiplets from the same datasets. Multiplet class statistical enrichment was determined as described above. Random multiplets of degree 2 were created by aggregating the read counts of randomly sampled cells. We determined the number of matched ligand-receptor pairs as follows. Using TP10K normalized scRNA data, we called a gene ‘expressed’ in a barcode if its normalized count was greater than the 25^th^ percentile of expression across all barcodes. For each barcode in a dataset, we counted the number of expressed ligands whose receptor was also expressed. We compared the distributions of the number of co-expressed ligand-receptor pairs across the three barcode classes (enriched multiplet types, non-enriched multiplet types, random multiplets) using Wilcoxon testing.

### Identifying cell-cell interactions in the small intestine

We obtained scRNA-seq data and metadata, including cell-type annotations, from Haber et al.(21) UMAP plots were drawn using Seurat(11). Neighbor-seq was run for n=10 iterations using default parameters. Cell-cell interactions were considered significant for those interactions with mean counts>10 and combined p-value<0.05. The validation analysis in known intestinal singlets multiplets was done using data from Andrews et al.(6), who also provided metadata and cell-type annotations. Neighbor-seq was run with default parameters as above on the small intestine scRNA-seq data. The resulting classifier was used to predict interactions in the accompanying small intestine partially dissociated clumps. Interactions were kept for those with counts>10 and adjusted p-value<0.05. Quantitative comparison of cell-cell networks from different studies was done by converting the networks into adjacency matrices and computing the inner product correlation. A permutation-based test was used to calculate the statistical significance of the inner product correlation. For a pair of matrices, random symmetric adjacency matrices are generated with the same number of nodes and edges as the test matrix, and a distribution of inner product correlations is computed. The center of this distribution is compared to the actual inner product correlation between the two test matrices using a one-sided Wilcoxon test.

### Identifying cell-cell interactions in the lung

We obtained scRNA-seq data and metadata, including cell-type annotations, from Travaglini et al.(22) To reduce computational complexity, cell-type annotations were collapsed into the following parent classes: alveolar, basal, ciliated, club, endothelial, fibroblast, goblet, immune, mucous, smooth muscle. UMAP plots were drawn using Seurat(11). Neighbor-seq was run for n=10 iterations using default parameters. Cell-cell interactions were considered significant for those interactions with mean counts>10 and combined p-value<0.05. The validation analysis in known lung singlets and multiplets was done using data from Andrews et al.(6), who also provided metadata and cell-type annotations. Neighbor-seq was run with default parameters as above on the lung scRNA-seq data. The resulting classifier was used to predict interactions in the accompanying lung partially dissociated clumps. Interactions were kept for those with counts>10 and adjusted p-value<0.05. Quantitative comparison of cell-cell networks from different studies was done with the permutation-based inner correlation test as described for the small intestine analysis.

### Identifying cell-cell interactions in the spleen

We obtained scRNA-seq data and metadata, including cell-type annotations, from Madissoon et al.(23) To reduce computational complexity, cell-type annotations were collapsed into the following parent classes: CD34 progenitor, CD4 T-cells, CD8 T-cells, Cycling T-cells, dendritic cells, follicular B cells, germinal center B cells, innate lymphoid cells, macrophage, mantle B cells, monocytes, natural killer cells, plasma B cells, and platelets. UMAP plots were drawn using Seurat(11). Neighbor-seq was run for n=10 iterations using default parameters. This study contained data from 19 human samples. Cell-cell interactions were considered significant for those interactions found in >1 sample and with mean counts>5 and combined p-value<0.05. The validation analysis was done using spleen data from Tabula Muris(24), who also provided metadata and cell-type annotations. Cell-type annotations were appended with their cluster label, resulting in multiple B-cell, T-cell, and myeloid clusters. Neighbor-seq was run with default parameters as above. Cell-cell interactions were considered significant for those interactions with mean counts>5 and combined p-value<0.05.

### Identifying cell-cell interactions in pancreatic cancer

We obtained scRNA-seq data and metadata, including cell-type annotations, from Peng et al.(25) UMAP plots were drawn using Seurat(11). Neighbor-seq was run for n=10 iterations using default parameters. Cell-cell interactions were considered significant for those interactions with mean counts>10 and combined p-value<0.05. Ligand-receptor analysis was done as described above. Betweenness centrality was calculated using igraph(14). Hierarchical clustering of betweenness centralities scaled by cell-type was done using the pheatmap R package (https://CRAN.R-project.org/package=pheatmap). Spatial transcriptomic data for PDAC tumors was obtained from Moncada et al.(26) Cell-type scores for each spatial barcode were calculated as follows. We used the FindAllMarkers function from Seurat(11) to find differentially expressed genes for the PDAC cell-types in Peng et al(25). Only highly cell-type specific genes were found by using these parameters: logfc.threshold=2, min.diff.pct=0.5, min.pct=0.5. Spatial barcodes were TP10K normalized, and cell-type scores were calculated as the average expression of each cell-type gene signature. Scores Spearman correlations were computed using the cor.test function in R. Quantitative comparison of cell-cell networks from different studies was done by converting the networks into adjacency matrices and computing the inner product correlation. The Spearman correlation matrix of the transcriptomic data was used as the adjacency matrix, only keeping those edges with values r>0.2 and p<0.05. A permutation-based test was used to calculate the statistical significance of the inner product correlation as described above.

### Identifying cell-cell interactions in skin cancer

We obtained scRNA-seq data and metadata, including cell-type annotations, from Ji et al. (27). To reduce computational complexity, immune cell-types annotations were collapsed into parent lymphoid and myeloid classes. UMAP plots were drawn using Seurat (11). Neighbor-seq was run for n=10 iterations using default parameters. Cell-cell interactions were considered significant for those interactions with mean counts>10 and combined p-value<0.05. Ligand-receptor analysis was done as described above. Cell-type scores for each spatial barcode was calculated as follows. We used the FindAllMarkers function from Seurat (11) to find differentially expressed genes for the cell-types in Ji et al. Only highly cell-type specific genes were found by using these parameters: logfc.threshold=0.5, min.diff.pct=0.52. Marker genes were only kept if they were differentially expressed in <5 cell types. Spatial barcodes were TP10K normalized, and cell-type scores were calculated as the average expression of each cell-type gene signature. Scores Pearson correlations were computed using the cor.test function in R.

## RESULTS

Neighbor-seq identifies physical cell interactions by using a machine learning classifier that annotates the cellular composition of each scRNA-seq barcode. Briefly, scRNA-seq data is filtered to remove low quality cells, the remaining barcodes are clustered based on their gene expression, and cluster marker genes are identified using a standard approach (11). A vast majority of the barcodes typically represent genuine single cells, with a minority being doublets or multiplets. Next, a training set of artificial multiplets is constructed representing all possible multiplet types by randomly sampling cells from each cell cluster and aggregating their raw read counts. The artificial multiplets can be homotypic (same cell-type) or heterotypic (different cell-type) and can be of order 2 (e.g. AA, AB, BB etc.) or higher (e.g. AAA, ABA, etc.). A Random Forest cell-type classifier is trained based on the expression of marker gene sets from *n* (default: 100) randomly sampled barcodes from each artificial multiplet type, as well as *n* barcodes from each original cell type cluster, which are predominantly singlets. Neighbor-seq then applies the classifier to annotate each barcode in the original dataset and identifies those classified as multiplets. An ensemble annotation result is generated by iterating the algorithm. Neighbor-seq computes the enrichment and corresponding statistical significance of the observed frequencies of multiplet types compared to that expected based on the prevalence of the original cell-types in the dataset. Accordingly, Neighbor-seq constructs a network representing significant cell-type interactions and enables transcriptomic analysis of interacting cells.

### Benchmarking and validation with known multiplets

We evaluated Neighbor-seq’s ability to identify physically interacting cells via (1) annotation of multiplets with known composition, (2) detection of known doublets in diverse tissues, and (3) identification of known cellular architectures in solid tissues. First, we tested Neighbor-seq’s ability to annotate barcode composition in a controlled setting. We obtained scRNA-seq data (6) for three cancer cell lines that had been sorted as either singlets or multiplets of known composition (**Fig. 1B**); we trained Neighbor-seq on the singlet data and used it to annotate singlet and multiplet barcode compositions, which contained between 1-4 cells in nonsymmetrical combinations. Neighbor-seq classified all droplets with high accuracy (AUC: singlets: 1, doublets: 0.99, triplets: 0.94, quadruplets: 0.82; **Fig. 1C**), indicating that it can reliably infer barcode composition.

Next, we benchmarked Neighbor-seq against 9 other published doublet-finding methods using 16 datasets of diverse tissue types in which doublets were experimentally annotated (10) (**Table S1**). Neighbor-seq performed better or equally well to all published methods in identifying doublets (Wilcoxon p > 0.8, **Fig. 1D**), but with the unique advantages of (1) determining multiplet cell-types, (2) identifying significantly enriched cell-type interactions, (3) requiring no parameter tuning, and (4) making no assumptions about the underlying doublet frequencies, which ranged from 2.5% to 37% in the benchmark datasets (**Fig. S1A)** and may not be known *a priori* in a real dataset. Moreover, Neighbor-seq is also the only method that can annotate higher order multiplets and can transfer knowledge across datasets.

We further hypothesized that since doublets contain neighboring cells, they would have increased co-expression of ligand-receptor pairs. Indeed, enriched doublets-types co-expressed significantly more ligand-receptor pairs than non-enriched doublets, and significantly more pairs than randomly synthesized doublets (both Wilcoxon p < 2.2e-16, **Fig. 1E, Fig. S1B**, see Methods for details). This was true in aggregate using two different ligand-receptor databases (15,16) and for each benchmark study individually (**Fig. S1B**).

Neighbor-seq performed exceptionally well at classifying singlets and all artificial doublet types (average AUC = 0.99, **Fig. 1F**) and performed significantly better than expected by chance when cell-type labels were randomized (cline_ch data; average AUC = 0.501, Wilcoxon p < 2.2e-16, **Fig. S1C-D**). Neighbor-seq classification accuracies across all benchmark datasets were similarly high (all AUC > 0.92, **Fig. S1E**). While Neighbor-seq is the only method designed to infer doublet-types, scDblFinder also proposes doublet compositions, but when comparing the stability of doublet-type counts across multiple runs on the same dataset, Neighbor-seq results were significantly more stable (Wilcoxon p < 2.2e-16, **Fig. 1G**). These results indicate that Neighbor-seq successfully and identifies doublets across a range of tissue types and reproducibly annotates their cellular composition. Taken together, these attributes make Neighbor-seq first in the class of innovative computational methods that can help reconstruct physical cell interaction networks from single-cell sequencing data.

### Identifying cell-cell interactions in the small intestine

We examined whether Neighbor-seq could identify known interaction architectures in multiple tissue types with varying levels of cell-type diversity and organization. First, we tested Neighbor-seq on a scRNA-seq survey of the small intestinal epithelium containing 11,666 cells from n=2 mice (21). The small intestine consists of alternating units of villi and crypts. Paneth cells in the crypt protect the residing stem cells, which differentiate and migrate upwards through various progenitor stages until becoming mature enterocytes, while goblet cells are scattered throughout villus (28) (**Fig. 2A**). We identified the major cell types (**Fig. 2B**) and used Neighbor-seq to recover their cellular adjacencies. Neighbor-seq correctly detected Paneth-stem interactions, a progression from stem to mature enterocytes, and multiple goblet cell interactions in the villus (**Fig. 2C**, see Methods for details). We confirmed this microanatomy by training Neighbor-seq on a separate study of 5,279 small intestinal singlets and used it to identify interactions in a dataset of 3,671 intestinal multiplets (6) (**Fig. 2D**). This validation analysis revealed a similar interaction network as obtained only from scRNA-seq data (**Fig. 2E**). Differences between the schematic illustration and the two networks are primarily due to different cell types annotated in the two datasets and different optimized network layouts; nonetheless, both networks recapitulate a progression from the Paneth-stem crypt to the mature enterocytes at the top of the villus whilst passing through transition and progenitor cells. Furthermore, both networks supported known ligand-receptor signaling (28), such as *LGR, LRP, BMP*, and *NOTCH*-mediated interactions between Paneth and stem cells. The directionality of this signaling could be inferred by inspecting the stem and Paneth cell singlets data, in which stem cells had significantly higher expression of the aforementioned genes compared to Paneth cells (Wilcoxon testing, p<2e-16 for all four genes). To quantitatively compare the networks obtained from the two studies, we harmonized the cell labels based on cell ontology and represented them as adjacency matrices (**Fig. 2F)**. The two matrices were significantly more correlated than expected by chance (inner product correlation = 0.53, Wilcoxon permutation test, p=4e-17, see Methods), indicating that Neighbor-seq identified similar cell networks from independent studies of the same tissue types.

**Figure 2.**
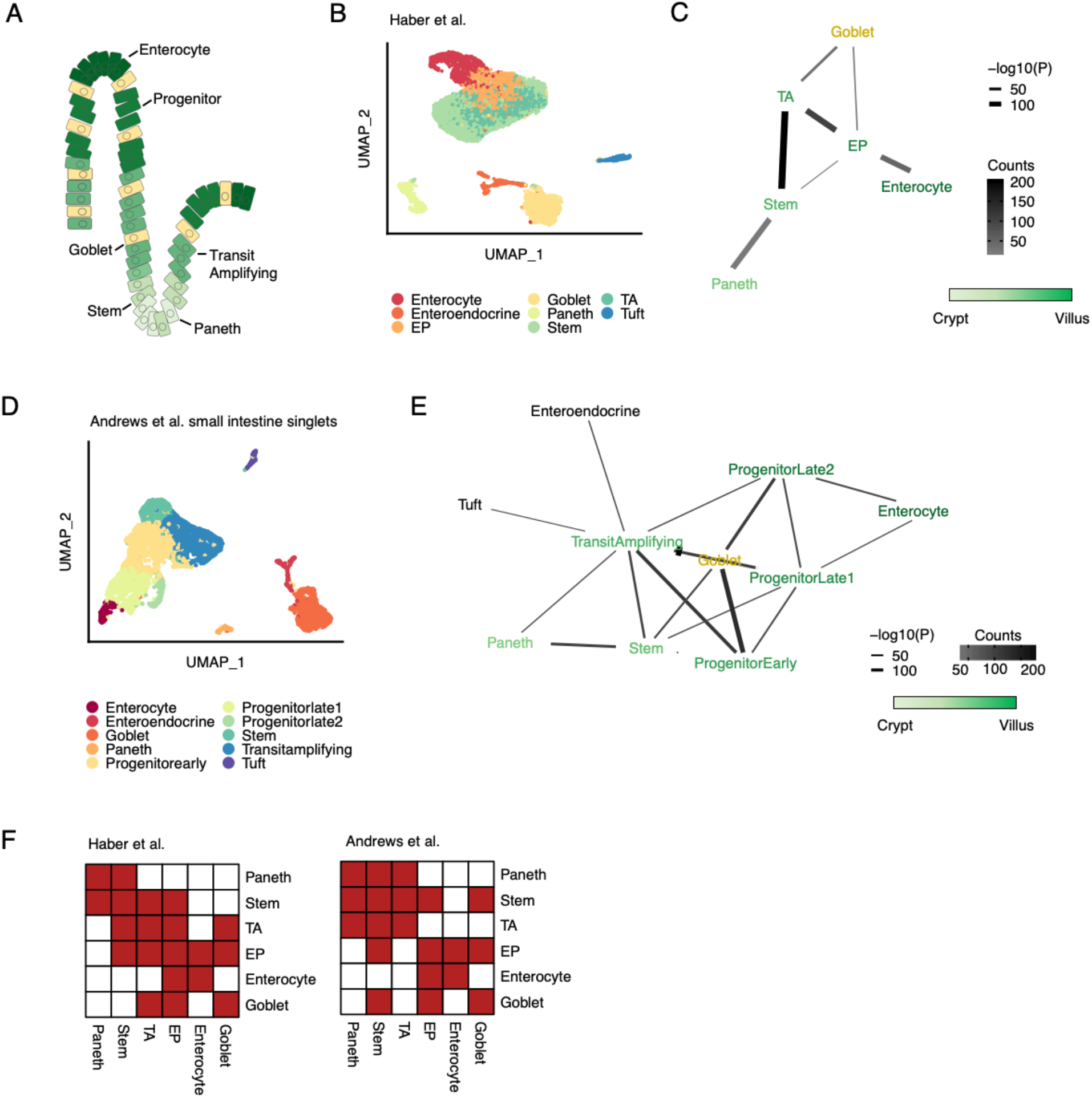
Detecting known microanatomical features of the small intestinal epithelium. **(A)** Illustration of the main cell types in the small intestinal crypt and villus. **(B)** Uniform manifold approximation and projection (UMAP) of 11,665 small intestinal cells (21) from the duodenum, jejunum, and ileum (n=2 mice) colored by cell type. TA, transit amplifying; EP, enterocyte progenitor. **(C)** Network diagram of significant cell type interactions from (B) identified by Neighbor-seq. Neighbor-seq is run for n=10 iterations, and interactions are shown for edges with a mean count > 10 and enrichment score combined adjusted p-value < 0.05 (see Methods). Edge thickness represents interaction p-value and edge color represents counts. Green color scale represents anatomical progression from crypt to villus. **(D)** UMAP of 5,279 cells from the small intestine (6) (n=1 mouse) colored by cell type. Neighbor-seq is trained on these cells and used to predict the interaction network in a dataset of partially dissociated intestinal clumps. **(E)** Network diagram of significant cell type interactions identified by Neighbor-seq from 3,671 small intestinal clumps. Methods, edge color and thickness, and colors scale are the same as in (C). **(F)** Adjacency matrix representation of the networks from (C) and (E). Cell labels were harmonized based on cell ontologies. Red color indicates the presence of a connection, white indicates no connection.

### Identifying cell-cell interactions in the lung

Second, we tested Neighbor-seq’s ability to identify microanatomical structures in the lung, which contains interactions between cells of multiple lineages. The terminal bronchioles contain ciliated epithelium and club cells, smooth muscle, mucous secreting cells, and basal stem cells; gas exchange occurs in the alveoli, which contain alveolar pneumocytes and macrophages and are lined by vessels, fibroblasts, and smooth muscle (22) (**Fig. 3A**). We analyzed a human lung cell atlas containing 65,662 cells from n=5 human donors (22), identified the cell-types (**Fig. 3B**) and recovered an interaction network (**Fig. 3C**, see Methods for details). Neighbor-seq correctly separated the bronchiolar and alveolar compartments and identified the gas exchange membrane between alveolar and endothelial cells. We confirmed detection of these cell-cell interactions in a dataset of lung singlets and multiplets (6) (**Fig. 3D**). Training Neighbor-seq on the singlet data and deconvoluting interactions from the known multiplets again revealed bronchiolar and alveolar compartments as well as several immune cell interactions (**Fig. 3E**). Neighbor-seq networks also corroborated known signaling pathways in lung development (29), such as by expression of *NOTCH, FGF*, and *EGF* ligands in basal-cell neighbors. Representing the networks derived from the two studies as adjacency matrices with harmonized cell-ontology labeling further indicated significant similarity in the cell-cell connections identified (**Fig. 3F**, inner product correlation=0.73, Wilcoxon permutation test, p=5e-17, see Methods).

**Figure 3.**
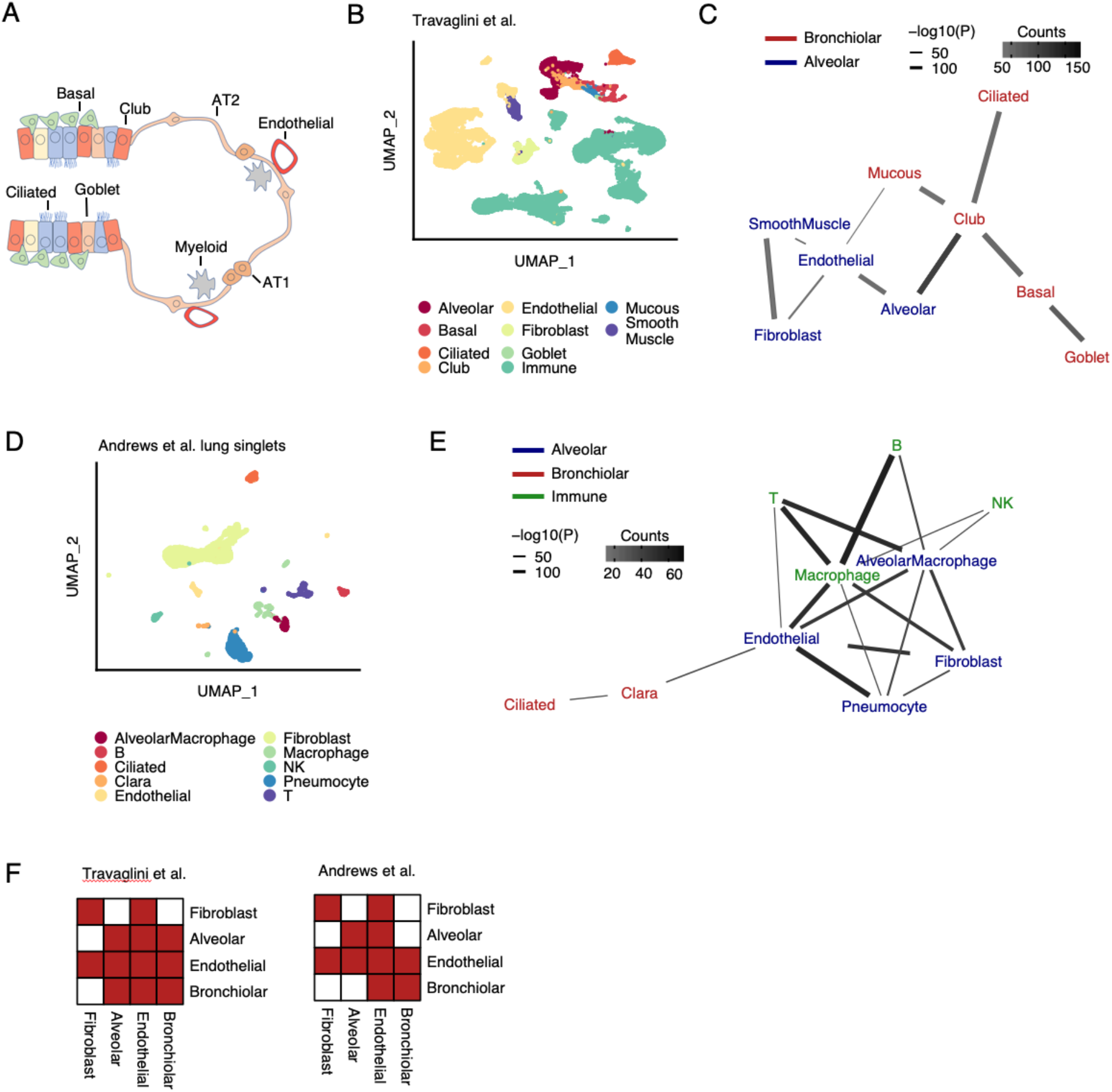
Detecting known microanatomical features of the terminal respiratory tract. **(A)** Illustration of the main cell types in the terminal bronchioles and alveolus. AT1, alveolar type 1 cell, AT2, alveolar type 2 cell **(B)** Uniform manifold approximation and projection (UMAP) 65,662 cells (21) from the terminal respiratory tract (n=5 human samples) colored by cell type. **(C)** Network diagram of significant cell type interactions from (B) identified by Neighbor-seq, colored by known microanatomical compartment. Data is shown for n=10 iterations, mean counts > 10, combined p-value < 0.05. **(D)** UMAP of 6,084 mouse lung cells (6) (n=3 mice) colored by cell type. Neighbor-seq is trained on these cells and used to predict the interaction network in a dataset of partially dissociated lung clumps. **(E)** Network diagram of significant cell type interactions identified by Neighbor-seq from 4,729 lung clumps. Methods, edge color and thickness, and colors scale are the same as in (C). **(F)** Adjacency matrix representation of the networks from (C) and (E). Cell labels were harmonized based on cell ontologies. Red color indicates the presence of a connection, white indicates no connection.

### Identifying cell-cell interactions in the spleen

Third, we used Neighbor-seq to predict interactions in the splenic white pulp, a structure not held by tight junctions but whose organization is chemokine driven (30). The white pulp is surrounded by myeloid cells and consists of separate B- and T-cell zones, with B-cell follicles further separating into germinal centers and mantle zones and T-cells interacting with antigen presenting myeloid cells (30) (**Fig. 4A**). We used Neighbor-seq to analyze a dataset of human spleen containing 94,050 immune cells from n=19 human samples (23) (**Fig. 4B**, see Methods for details). Neighbor-seq correctly identified the B and T-cell zones; germinal centers did not interact with non-B-cells, and myeloid cells interacted with both B and T lymphocytes (**Fig. 4C**). We confirmed these structures by analyzing splenic tissue from the Tabula Muris (24) containing 9,552 cells of lymphoid and myeloid origin (**Fig. 4D**). Neighbor-seq again identified B and T-cell zones with myeloid interactions (**Fig. 4E**). Collectively, these analyses confirmed that Neighbor-seq can correctly identify direct cell-cell interactions and microanatomies *in vivo* in diverse tissue types.

**Figure 4.**
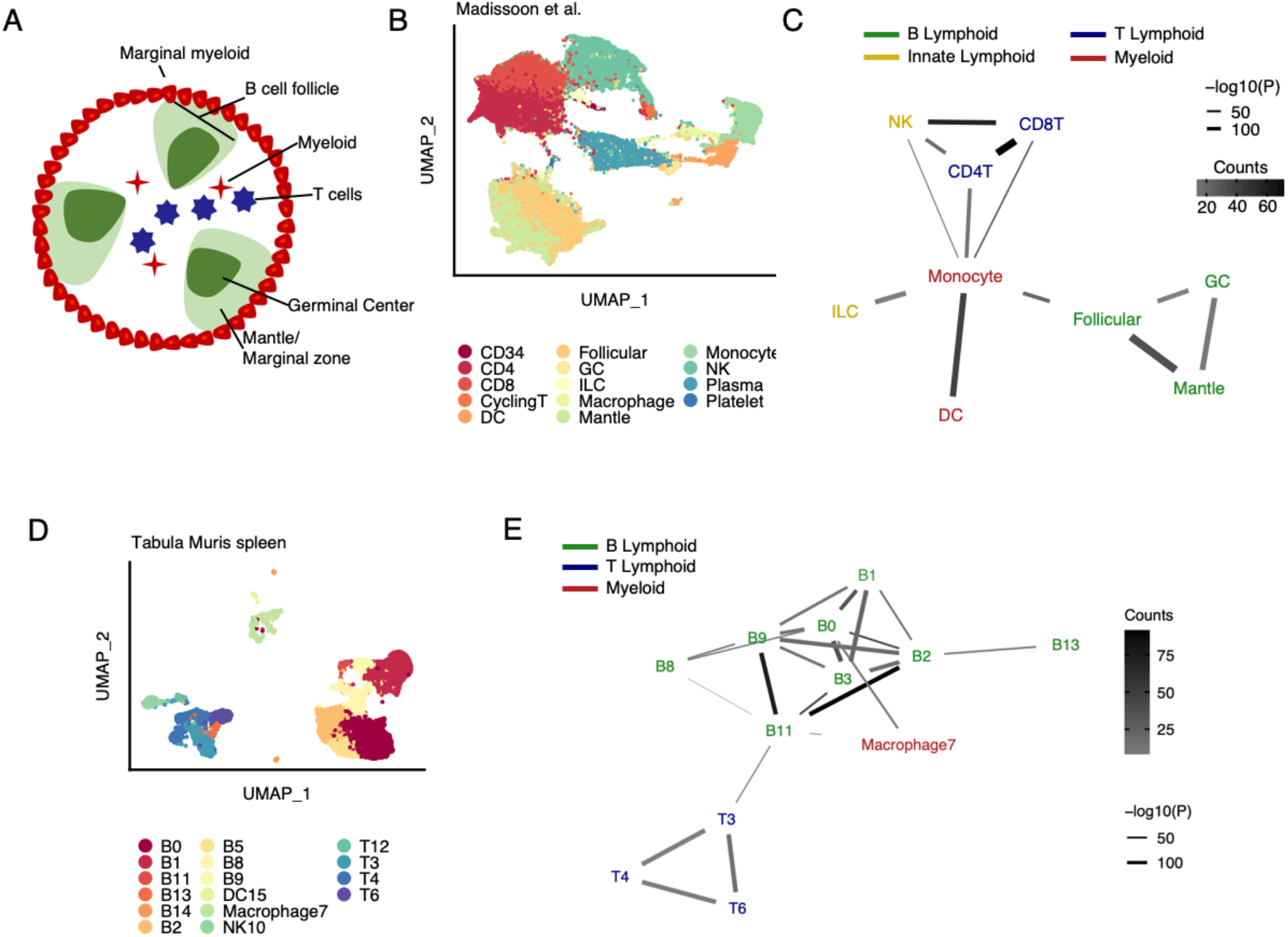
Detecting known microanatomical features of the splenic white pulp. **(A)** Illustration of the main cell types in the splenic white pulp. **(B)** Uniform manifold approximation and projection (UMAP) of 94,050 splenic white pulp immune cells (Madissoon et al., 2019) (n=19 human samples) colored by cell type. DC, dendritic cell; GC, germinal center; ILC, innate lymphoid cell; NK, natural killer cell. **(C)** Network diagram of significant cell type interactions from (B) identified by Neighbor-seq, colored by known cell type lineage. Data is shown for n=10 iterations, mean counts > 5, combined p-value < 0.05. **(D)** UMAP of 9,552 murine immune cells from Tabula Muris (Schaum et al., 2018) colored by cell type. **(E)** Network diagram of significant cell type interactions from (D) identified by Neighbor-seq, colored by known cell compartment. Methods, edge color and thickness, and colors scale are the same as in (C).

### Identifying cell-cell interactions in pancreatic cancer

We next used Neighbor-seq to analyze inter-cellular interactions in pancreatic ductal adenocarcinoma (PDAC). We obtained scRNA-seq data for 24 tumors and 11 control samples (25); these tissues had normal to poorly differentiated histopathology, were from anatomic locations throughout the pancreas (**Table S2**), and in total contained 57,530 cells from a mixture of normal and malignant epithelial cells and immune, stromal, and endothelial cells (**Fig. 5A**). Neighbor-seq detected 17,580 doublets (31% of barcodes), of which 1,373 were heterotypic doublets (2.3% of all barcodes) – this fraction of heterotypic cell neighbors was typical for all datasets analyzed.

**Figure 5.**
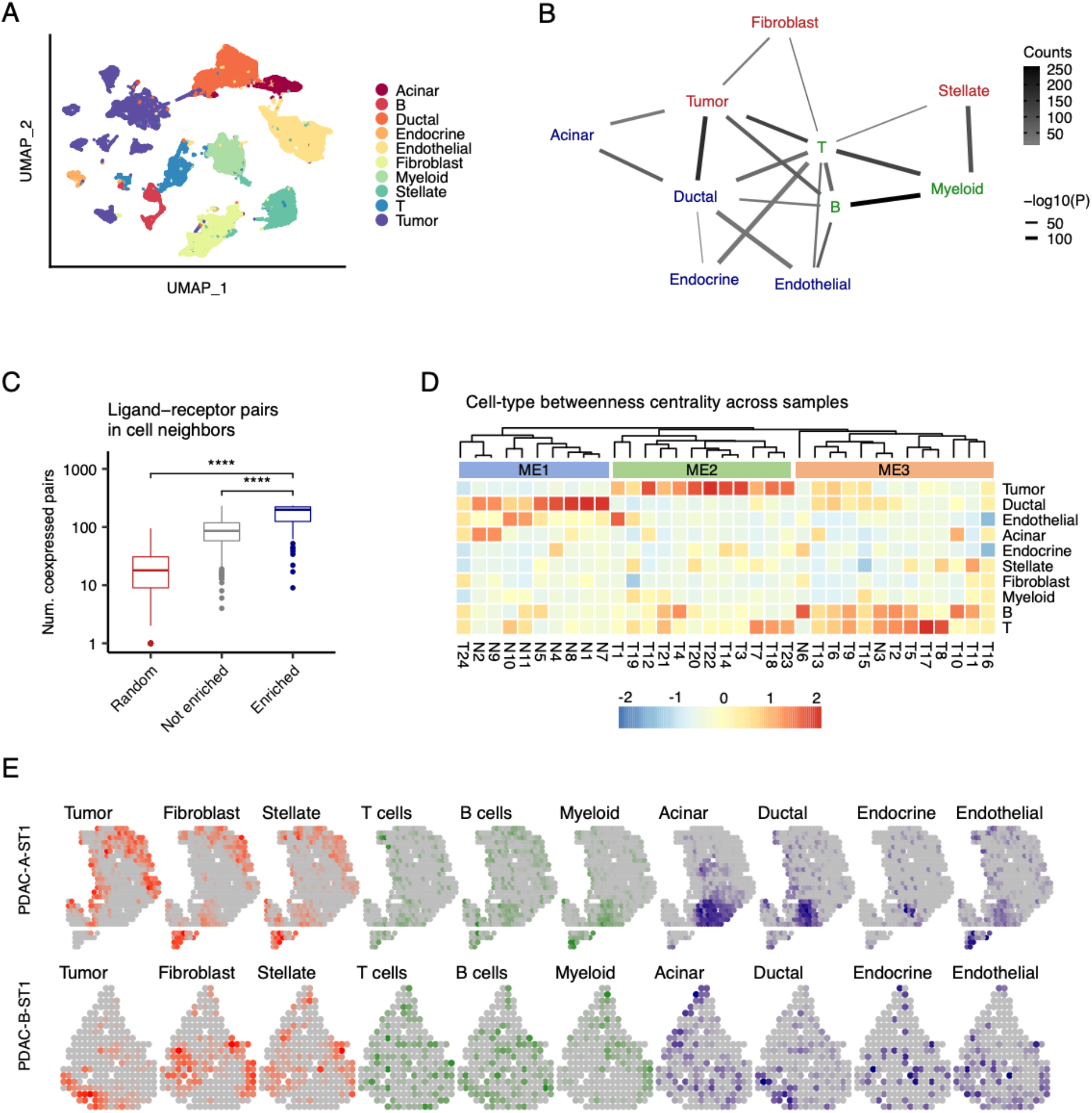
Identification of cell type interactions in pancreatic cancer. **(A)** Uniform manifold approximation and projection (UMAP) of 57,530 primary human cells (Peng et al., 2019) from n=24 pancreatic ductal adenocarcinomas and n=11 control pancreatic tissues colored by cell type. **(B)** Network diagram of significant cell type interactions from (A) identified by Neighbor-seq, colored by known cell compartment. Edge thickness represents interaction p-value and edge color represents counts. Neighbor-seq is run for n=10 iterations, and interactions are shown for edges with a mean count > 10 and enrichment score combined adjusted p-value < 0.05 (see Methods). **(C)** Comparison of the number of co-expressed ligand-receptor pairs in enriched (statistically significant) doublet types, not enriched doublets, and random synthetic doublets from (A). Boxplots show median (line), 25^th^ and 75^th^ percentiles (box) and 1.5xIQR (whiskers). Points represent outliers. Wilcoxon tests, ****p<0.0001. **(D)** Heatmap and hierarchical clustering of cell type betweenness centrality in interaction networks for each sample. Rows represent cell types and columns represent samples; T#, tumor sample; N#, normal sample; ME#, microenvironment. **(E)** Spatial transcriptomic maps of two pancreatic tumors (Moncada et al., 2020a) with n=428 (top) and n=224 (bottom) barcodes, colored by cell-type abundance scores. Correlation of abundance score of multiple cell type pairs, identified in Fig 3B, were significant (tumor and ductal cells: A: r=0.19, p=2e-2; B: r=0.44, p=4e-10; tumor cells and fibroblasts: A: r=0.30, p=8e-9; ductal and acinar cells: A: r=0.16, p=2e-2; ductal and endocrine cells: A: r=0.29, p=5e-8; ductal and endothelial cells A: r=0.41, p=1e-17; Spearman correlation). See **Fig. S3** for additional instances.

Out of 45 possible heterotypic cell-type pairs, Neighbor-seq identified 19 enriched interactions (**Fig. 5B**). Normal pancreatic epithelium (ductal, endocrine, acinar cells) interacted with each other and with endothelial cells, whereas tumor cells had stronger connections with ductal cells and fibroblasts and immune cells interacted with most cell-types. These interactions were detected across multiple samples, the most common being B cell-myeloid, ductal-tumor, and tumor-T cell interactions (**Fig. 5B, Fig. S2A, Table S3**). Again, enriched cell neighbors co-expressed significantly more matched ligand-receptor pairs than non-enriched doublet-types or random synthetic doublets (Wilcoxon p < 2.2e-16, **Fig. 5C**). The detected neighboring cell communications were also consistent with existing literature. For example, *BTK-*dependent B-cell interactions with *FCγR+* myeloid cells were recently implicated in PDAC progression (31), and we observed significantly greater *BTK* (Wilcoxon, p=6.3e-40) and *FCGR1A* expression (Wilcoxon, p=6.8e-59) in B cell-myeloid doublets compared to all other doublet-types. This highlights how Neighbor-seq can be used to identify interaction-specific transcriptional changes with functional relevance.

The topology of cell-cell interactions varied across samples, which could be due to intra- or inter-tumor heterogeneity in cell interactions or differential presence/absence of doublet-types in a sample. We computed the betweenness centrality of cell-types in each sample, and clustering of these centrality scores revealed three classes of tissue microenvironments (ME, **Fig. 5D**). ME1 contained mostly normal ductal epithelial connections, ME2 was dominated by tumor cell connections, and ME3 by immune cell edges (**Fig. 5D**). These ME classes had significantly different histopathological characteristics (Fisher’s exact test; pathology: p=5e-4; stage: p=5e-4; lymph node invasion: p=3e-3). These analyses indicate that Neighbor-seq can uncover direct cellular interactions and tumor microenvironment characteristics with functional or clinical relevance.

Next, we analyzed spatial transcriptomic data for pancreatic ductal adenocarcinoma (PDAC) samples from a published study (26) where there were 243-996 spatially annotated barcodes per sample, each capturing the aggregated transcriptomic makeup of approximately 20-70 cells in tumor and adjacent normal tissue microenvironments. We observed organizational contexts similar to what we identified from scRNA-seq data alone. We used marker genes to calculate cell-type scores for each spatial barcode (see Methods; PDAC-A: n=428, PDAC-B: n=224 spatial barcodes) and visualized the distribution of cell-types in the tissues (**Fig. 5E**). Tumor cells co-localized with ductal cells (Spearman correlation; A: r=0.19, p=2e-2; B: r=0.44, p=4e-10) and fibroblasts (Spearman correlation; A: r=0.30, p=8e-9). Ductal cells co-localized with acinar cells (Spearman correlation; A: r=0.16, p=2e-2), endocrine cells (Spearman correlation; A: r=0.29, p=5e-8), and endothelial cells (Spearman correlation; A: r=0.41, p=1e-17). B and T cells were scattered throughout the tissue, but myeloid cells did not significantly co-localize with tumor cells. We observed similar patterns when visualizing spatial transcriptomic data for n=7 other PDAC tumors (**Fig. S3A**). These findings are broadly consistent with the architecture of cell-networks we deduced from scRNA-seq data of a different cohort of PDAC tumors (**Fig. 5C**), suggesting that despite intra- and inter-tumor heterogeneity, the topology of cell-cell interactions may be similar. We represented the cell-cell networks derived from scRNA-seq and the spatial transcriptomic data as cell adjacency matrices (**Fig. S2B**, see Methods). Inner product correlation further indicated that the cell adjacencies identified from the two studies were significantly more similar than expected by chance (r=0.33, p=2e-11). This validation across two different technology platforms reinforces Neighbor-seq’s ability to infer cellular connectomes from scRNA-seq data and bridge between single-cell and spatial transcriptomics, which has a resolution of 10^1^-10^2^ cells.

### Identifying cell-cell interactions in skin cancer

Lastly, we used Neighbor-seq to analyze physical cell interactions in cutaneous squamous cell carcinoma (SCC). We obtained scRNA-seq data for 10 tumors, of which 4 tumors also had spatial transcriptomic data (27). These tissues had moderate or well differentiated histology, were from various primary sites on the body, and the dataset in total contained 48,164 single-cell transcriptomes from 16 cell subtypes of epithelial, stromal, and immune origin (**Fig. 6A**). Neighbor-seq detected 15,626 doublets (32% of all barcodes), of which 5,346 were heterotypic doublets (11% of all barcodes). Of 105 possible heterotypic cell-type pairs, Neighbor-seq identified 38 significant pairwise interactions (**Fig. 6B**). Tumor cell populations were on the edge of the cell interaction network and showed significant interactions mostly with normal epithelium. Unlike in pancreatic cancer (**Fig. 5B**), immune cells were located peripherally in the network and interacted mostly with normal epithelium. We did not observe direct connections between lymphoid cells and any tumor subpopulation (**Fig. 6B**), which might suggest potential immune evasion.

**Figure 6.**
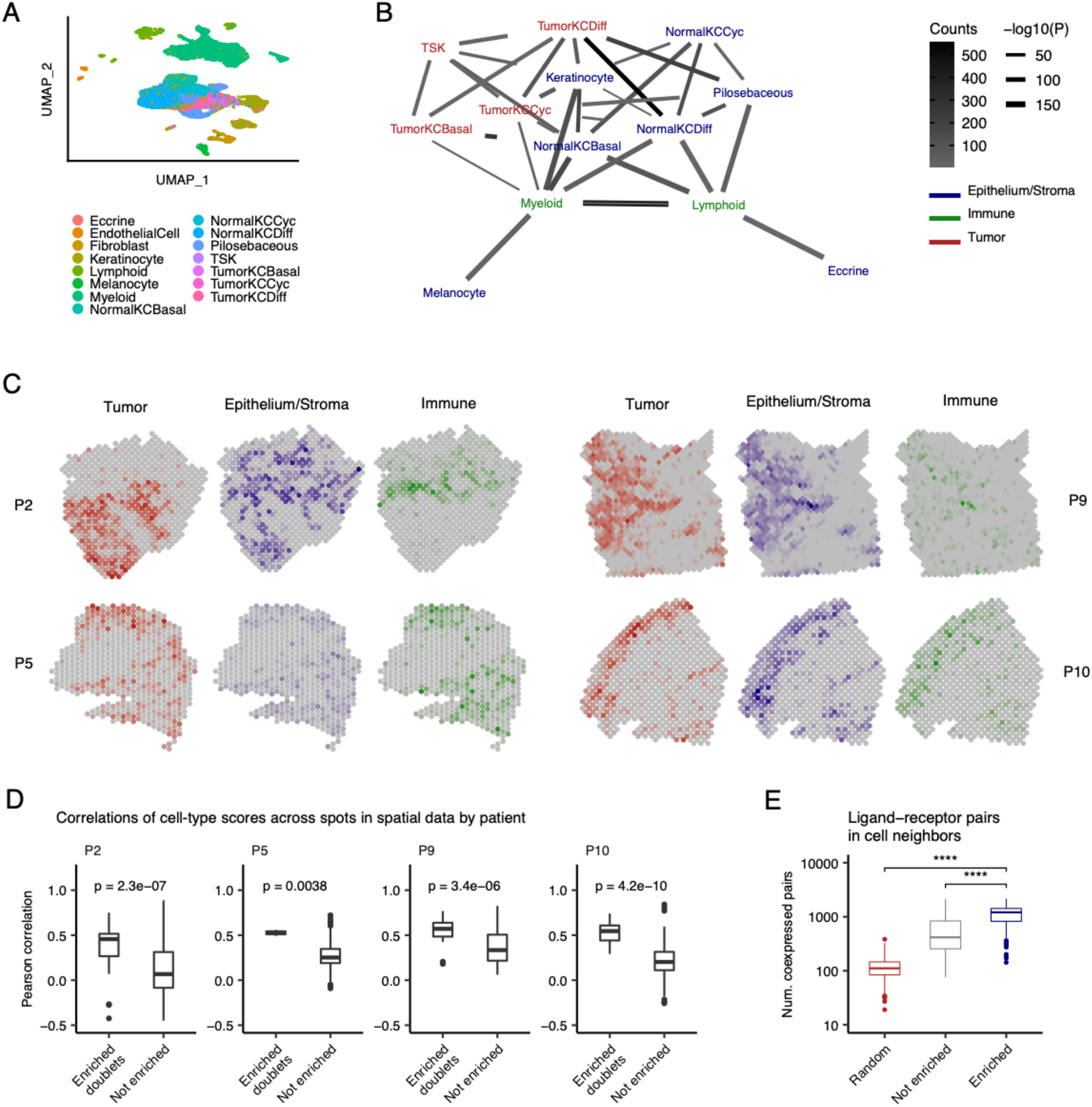
Identification of cell type interactions in skin cancer. **(A)** Uniform manifold approximation and projection (UMAP) of 48,164 primary human cells from n=10 cutaneous squamous cell carcinomas (26) colored by cell type. TSK, tumor-specific keratinocyte; KC, keratinocyte; Cyc, cycling; Diff, differentiating. **(B)** Network diagram of significant cell type interactions from (A) identified by Neighbor-seq, colored by known cell compartment. Edge thickness represents interaction p-value and edge color represents counts. Neighbor-seq is run for n=10 iterations, and interactions are shown for edges with a mean count > 10 and enrichment score combined adjusted p-value < 0.05 (see Methods). **(C)** Spatial transcriptomic maps of four patients (P2: n=666, P5: n=521, P9: n=1145, P10: n=608 barcodes) colored by cell-type abundance scores (see Methods). **(D)** Boxplots comparing spatial barcode cell-type abundance scores between cell-type pairs with enriched doublets from (B) compared to all other possible pairs. Boxplots show median (line), 25^th^ and 75^th^ percentiles (box) and 1.5xIQR (whiskers). Points represent outliers. Wilcoxon testing. **(E)** Comparison of the number of co-expressed ligand-receptor pairs in enriched (statistically significant) doublet types, not enriched doublets, and random synthetic doublets from (A). Boxplots are the same as in (D).

These general patterns identified from the inferred cell-interaction network agreed with observations made from spatial transcriptomic sequencing data of the same tumors profiled at 8,179 spots. Using the single-cell data, we identified cell-type marker genes and used these to calculate cell-type scores for each spatial sequencing spot in 4 patients (see Methods). Coloring the spatial maps by composite tumor, normal epithelium and stroma, or immune scores revealed spatial contexts similar to those we identified with Neighbor-seq from the single-cell data alone (**Fig. 6C**). Most prominently in patient P2, tumor populations occupied a separate region of the spatial map with minimal immune presence, while the adjacent regions were rich in normal epithelium, stromal, and immune genetic activity. We next compared the specific cell-type pair associations identified by Neighbor-seq (**Fig. 6B**) with the cell-type score correlations in the matched spatial data. For all patients, cell-types with enriched neighbor interactions in scRNA-seq were significantly more correlated with each other across spatial spots than were cell-type pairs that did not have enriched doublets in scRNA-seq (P2: p=2.3e-7, P5: p=3.8e-4, P9: p=3.4e-6, P10: p=4.2e-10; Wilcoxon; **Fig. 6D**). Lastly, we confirmed that the enriched cell neighbors in scRNA-seq exhibited increased pairwise crosstalk by assessing the number of co-expressed ligand-receptor pairs (see Methods). Consistent with our previous analyses, enriched doublets expressed significantly more ligand-receptor pairs than non-enriched doublets or randomly synthesized doublets (both Wilcoxon p<2.2e-16, **Fig. 6E**). Taken together, these results indicate that Neighbor-seq accurately identifies and annotates multiplets that contain interacting cells across a range of normal and diseased tissues, and these interactions are consistent with spatial contexts identified from both the same and different samples.

## DISCUSSION

In summary, Neighbor-seq infers direct cell-cell interactions by identifying undissociated cell multiplets in standard scRNA-seq data and classifying them according to their constituent cell-types, ultimately building the cellular interactome in diverse normal and diseased tissues. It shows high accuracy and reproducibility, and the results are in agreement with prior knowledge about tissue microenvironments in well-studied tissues and with spatial transcriptomic data. Neighbor-seq complements emerging methods (5–8) that are optimized for deconvoluting the transcriptomics of single cells and cell clumps. It, however, reconstructs tissue-scale cell interaction networks using undissociated multiplets from standard scRNA-seq alone, thereby eliminating the need for specialized sample preparation and boosting scalability. Neighbor-seq provides a framework to study the topology of cell-cell interactome leading to the organization of tissue microenvironment, bridging the gap in resolution between single-cell and spatial transcriptomics. Neighbor-seq is available at https://github.com/sjdlabgroup/Neighborseq.

## DATA AVAILABILITY

The Neighbor-seq resource and user-friendly documentations are freely available on Github at https://github.com/sjdlabgroup/Neighborseq

## SUPPLEMENTARY INFORMATION

**Figure S1**. Neighbor-seq performance on benchmark datasets.

**Figure S2**. Interactions in pancreatic cancer.

**Figure S3**. PDAC spatial transcriptomic maps colored by cell-type scores.

**Figure S4**. Overview of Neighbor-seq algorithm

**Figure S5**. Benchmarking the fast implementation of Neighbor-seq

**Table S1**. Summary of datasets analyzed in this study

**Table S2**. Clinical characteristics of PDAC patients and control samples; adapted from Peng et al., Nature Cell Research 2019

**Table S3**. Cell type interactions in pancreatic cancer

## ACKNOWLEDGMENTS

We acknowledge the Office of Advanced Research Computing (OARC) at Rutgers, The State University of New Jersey for providing access to the Amarel cluster URL: https://it.rutgers.edu/oarc.

## FIGURE CAPTIONS

**Figure S1. Neighbor-seq performance on benchmark datasets. (A)** Fraction and number of doublets found in benchmark datasets. **(B)** Comparison of the number of co-expressed ligand-receptor pairs in enriched (statistically significant) doublet types, not enriched doublets, and random synthetic doublets from the benchmarking studies in Fig. 1D using ligand-receptor data from CellTalkDB (16). Boxplots show median (line), 25^th^ and 75^th^ percentiles (box) and 1.5xIQR (whiskers). Points represent outliers. Wilcoxon tests, ****p<0.0001. (**C)** Comparison of the number of co-expressed ligand-receptor pairs in enriched (statistically significant) doublet types, not enriched doublets, and random synthetic doublets from benchmark datasets with experimentally annotated doubles (see **Table S1**). Boxplots are as in (B). **(D)** Neighbor-seq artificial multiplet classification performance on the cline-ch dataset for label-shuffled cell-types. AUC, area under the curve. **(E)** Comparison of distributions of area under the receiver operator curves (AUC) for artificial multiplet classification of the cline-ch dataset for true (**Fig. 1D)** vs. label-shuffled (**Fig. S1D**) data. Boxplots as in (A), Wilcoxon testing. **(F)** Average artificial multiplet type classification performance for all benchmark datasets. AUC, area under the receiver operator curve.

**Figure S2. Interactions in pancreatic cancer. (A)** Dot plot showing neighbor-types found across samples in n=24 pancreatic tumors and n=11 control pancreatic tissues (25). Rows represent neighbor-types and columns represent pancreas samples. T#, tumor sample; N#, normal sample. Neighbor-seq is run for n=10 iterations and data is shown for neighbor types with mean counts > 10 and combined adjusted p-value < 0.05. **(B)** Adjacency matrix representation of the networks from **Fig. 5B** and one tumor from **Fig. 5E**. Red color indicates the presence of a connection, white indicates no connection.

**Figure S3. PDAC spatial transcriptomic maps colored by cell-type scores. (A)** Spatial maps for n=7 pancreatic tumors (26) colored by cell-type scores (see Methods).

**Figure S4. Overview of Neighbor-seq algorithm**. Schematic diagram of key Neighbor-seq steps: (1) single-cell sequencing, cell type clustering, and marker gene identification, (2) enumerating neighbor-types, (3) random forest training on a dataset of artificial multiplets and singlets, (4) barcode composition prediction of the original dataset, (5) neighbor-type enrichment and cell-cell network construction.

**Figure S5. Benchmarking the fast implementation of Neighbor-seq. (A)** Benchmarking Neighbor-seq fast implementation doublet detection against 9 other methods using 16 datasets of diverse tissue types with experimentally annotated doublets (see Table S1). Comparison of singlet vs. doublet classification area under the receiver operator curve (AUROC) distributions. Boxplots show median (line), 25^th^ and 75^th^ percentiles (box) and 1.5xIQR (whiskers). Points represent outliers. **(B)** Neighbor-seq fast implementation barcode composition annotation performance of cell-line barcodes in **Fig. 1F**, plotted by known barcode type. AUC = area under the receiver operator curve.

