## Supplementary for "Reconstructing physical cell interaction networks from single-cell data using Neighbor-seq"

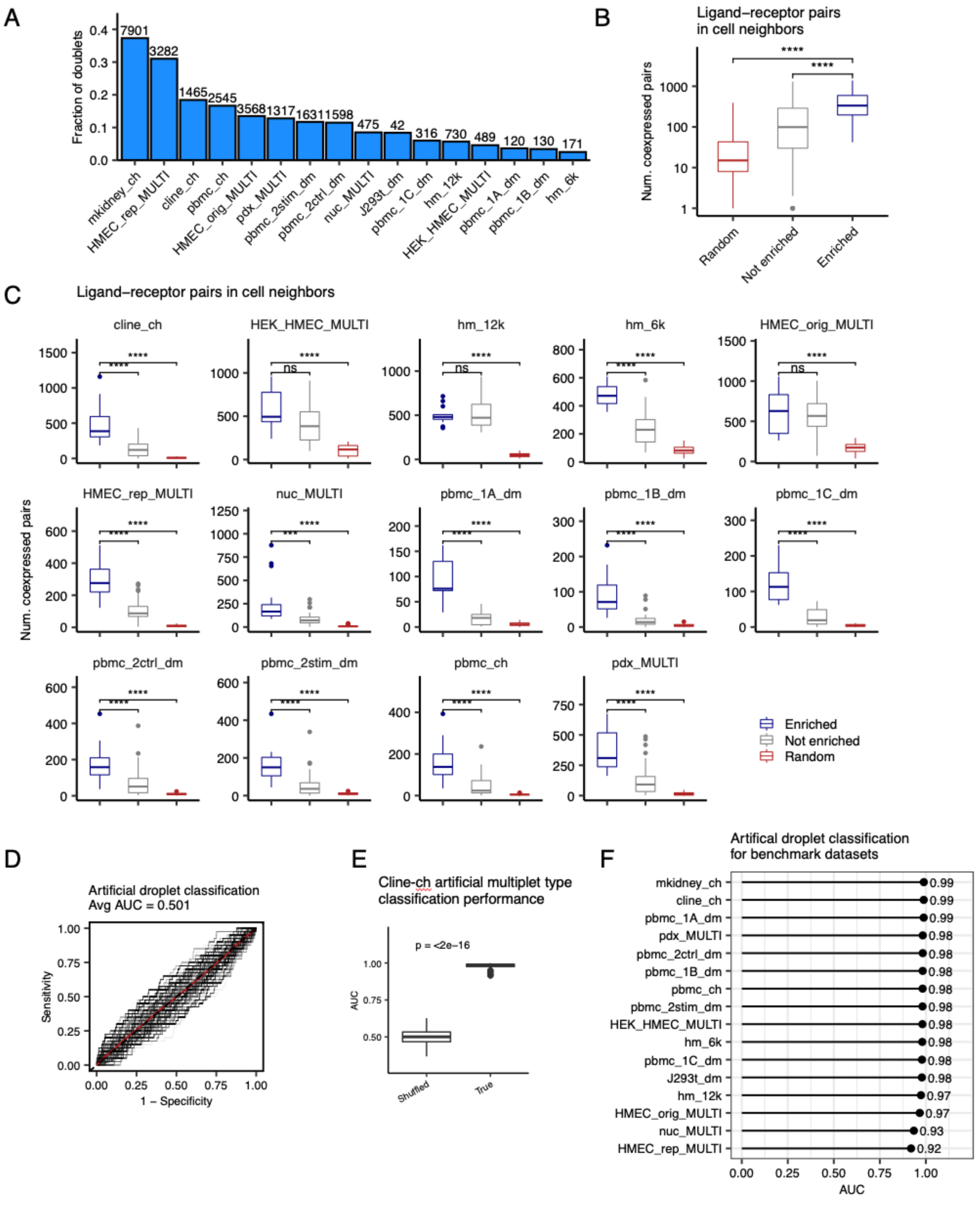


**Figure S1. Neighbor-seq performance on benchmark datasets**. **(A)** Fraction and number of doublets found in benchmark datasets. **(B)** Comparison of the number of co-expressed ligand-receptor pairs in enriched (statistically significant) doublet types, not enriched doublets, and random synthetic doublets from the benchmarking studies in Fig. 1D using ligand-receptor data from CellTalkDB (16). Boxplots show median (line), 25th and 75th percentiles (box) and 1.5xIQR (whiskers). Points represent outliers. Wilcoxon tests, ****$p<0.0001$. **(C)** Comparison of the number of co-expressed ligand-receptor pairs in enriched (statistically significant) doublet types, not enriched doublets, and random synthetic doublets from benchmark datasets with experimentally annotated doubles (see **Table S1**). Boxplots are as in (B). **(D)** Neighbor-seq artificial multiplet classification performance on the cline-ch dataset for label-shuffled cell-types. AUC, area under the curve. **(E)** Comparison of distributions of area under the receiver operator curves (AUC) for artificial multiplet classification of the cline-ch dataset for true (**Fig. 1D**) vs. label-shuffled (**Fig. S1D**) data. Boxplots as in (A), Wilcoxon testing. **(F)** Average artificial multiplet type classification performance for all benchmark datasets. AUC, area under the receiver operator curve.

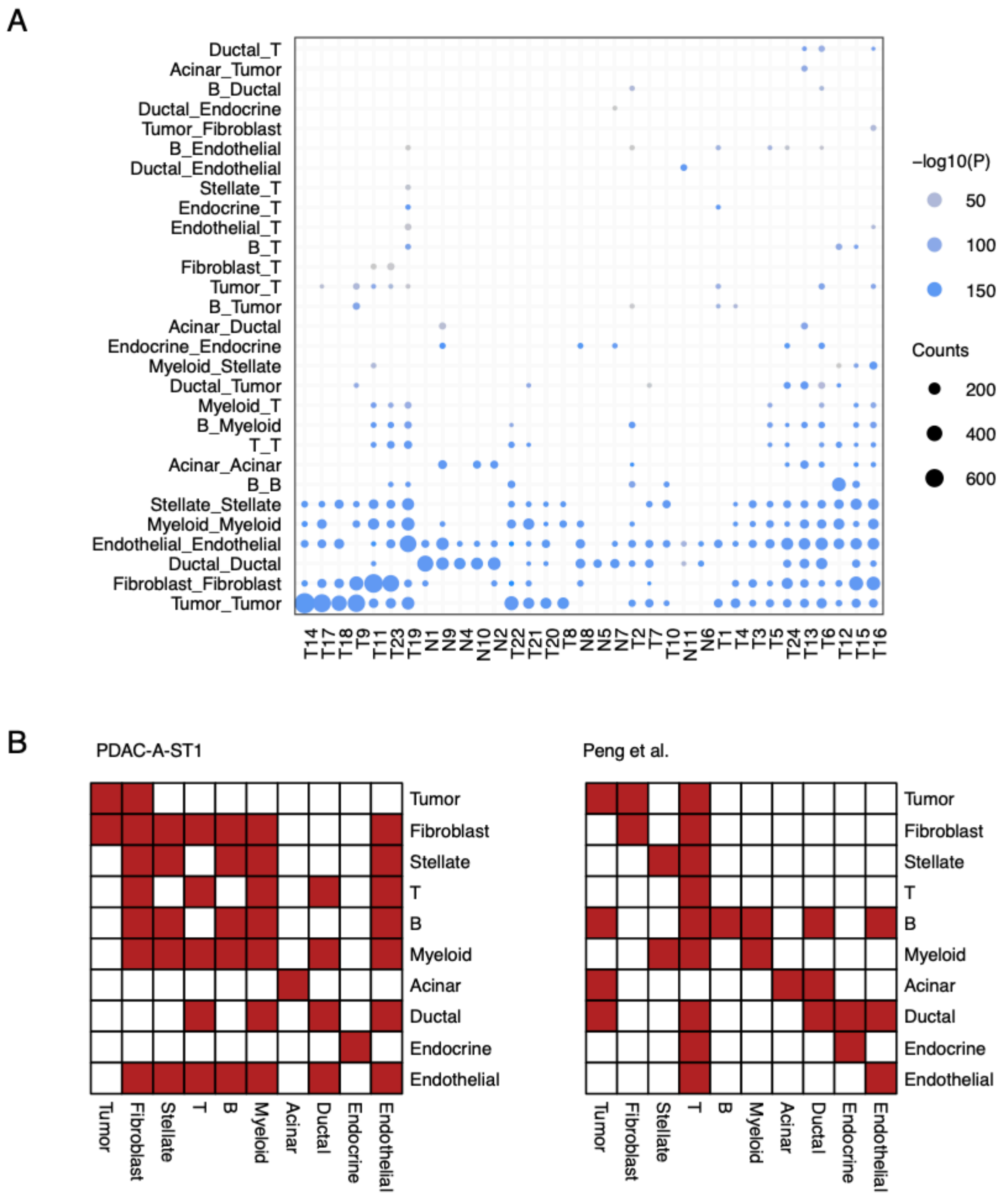


**Figure S2. Interactions in pancreatic cancer. (A)** Dot plot showing neighbor-types found across samples in n=24 pancreatic tumors and n=11 control pancreatic tissues (25). Rows represent neighbor-types and columns represent pancreas samples. T#, tumor sample; N#, normal sample. Neighbor-seq is run for n=10 iterations and data is shown for neighbor types with mean counts > 10 and combined adjusted p-value < 0.05. **(B)** Adjacency matrix representation of the networks from **Fig. 5B** and one tumor from **Fig. 5E**. Red color indicates the presence of a connection, white indicates no connection.

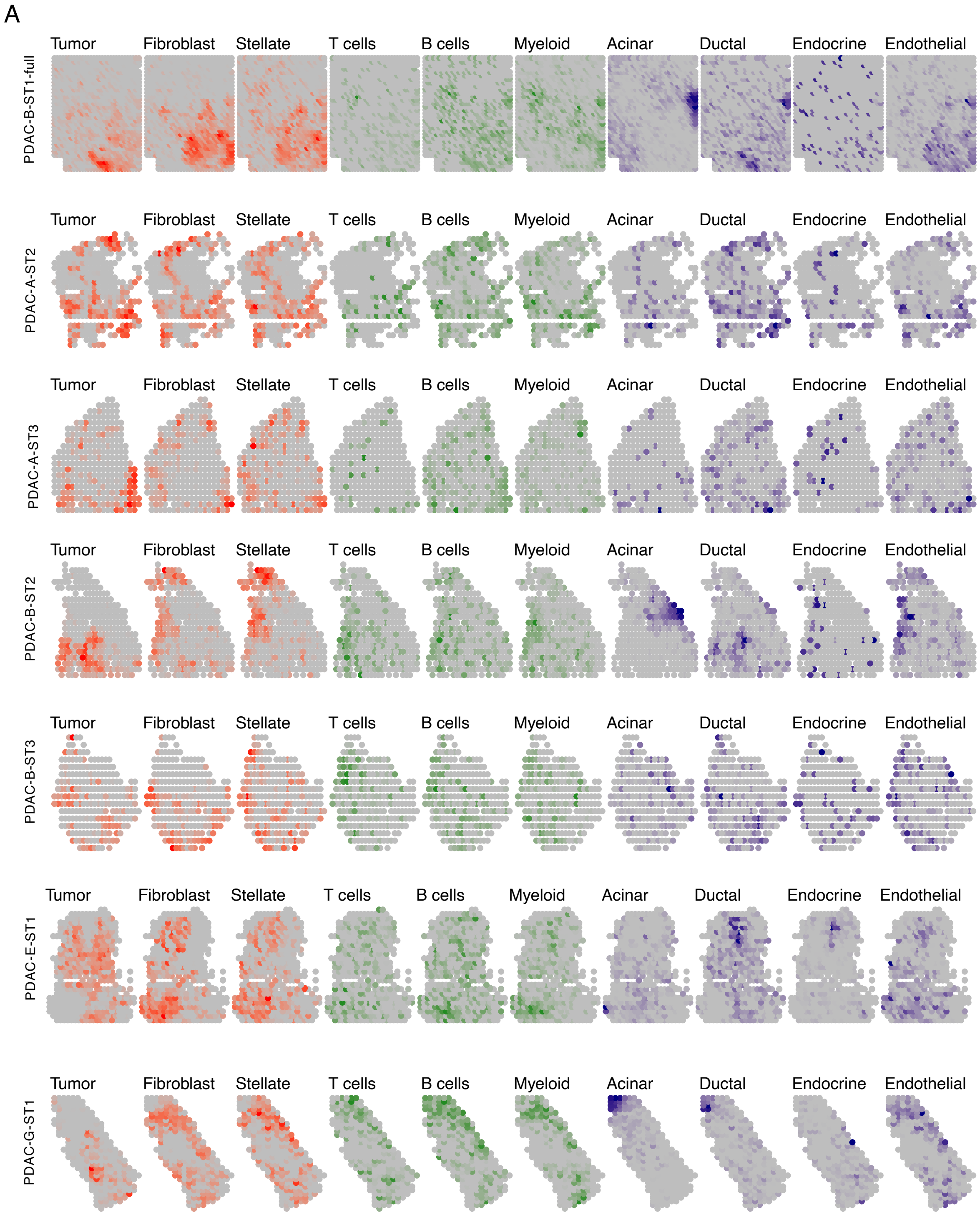


**Figure S3. PDAC spatial transcriptomic maps colored by cell-type scores. (A)** Spatial maps for n=7 pancreatic tumors (Moncada et al., 2020a) colored by cell-type scores (see Methods).

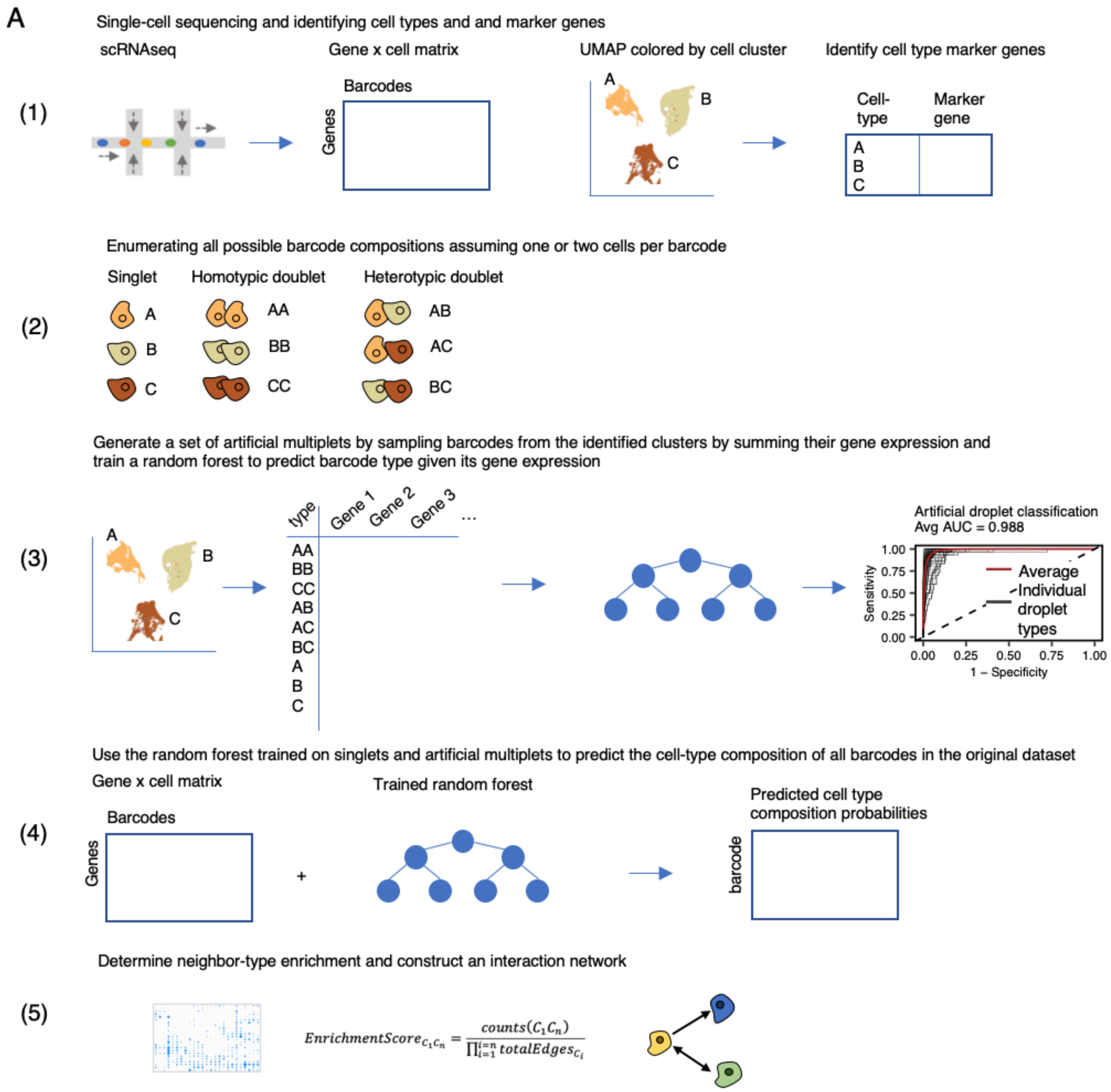


**Figure S4. Overview of Neighbor-seq algorithm.** Schematic diagram of key Neighbor-seq steps: (1) single-cell sequencing, cell type clustering, and marker gene identification, (2) enumerating neighbor-types, (3) random forest training on a dataset of artificial multiplets and singlets, (4) barcode composition prediction of the original dataset, (5) neighbor-type enrichment and cell-cell network construction.

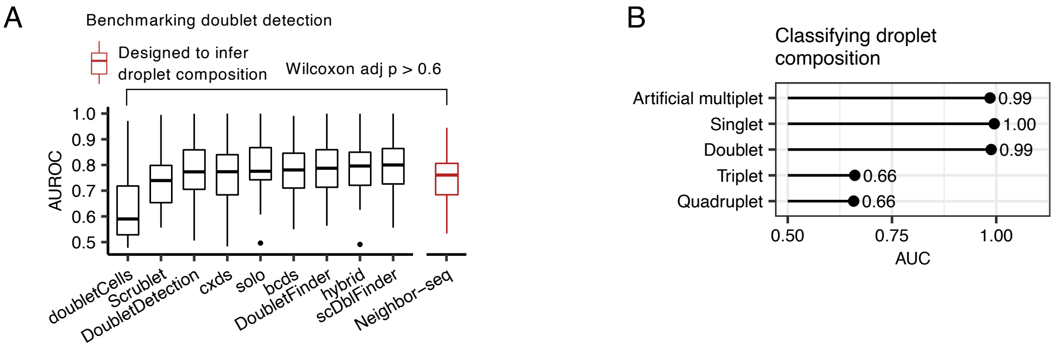


**Figure S5. Benchmarking the fast implementation of Neighbor-seq**. **(A)** Benchmarking Neighbor-seq fast implementation doublet detection against 9 other methods using 16 datasets of diverse tissue types with experimentally annotated doublets (see Table S1). Comparison of singlet vs. doublet classification area under the receiver operator curve (AUROC) distributions. Boxplots show median (line), 25$^{th}$ and 75$^{th}$ percentiles (box) and 1.5xIQR (whiskers). Points represent outliers. **(B)** Neighbor-seq fast implementation barcode composition annotation performance of cell-line barcodes in **Fig. 1F,** plotted by known barcode type. AUC = area under the receiver operator curve.
